# Rapid and Durable Protection Against Marburg Virus with a Single-Shot ChAd3-MARV GP Vaccine

**DOI:** 10.1101/2021.12.22.472410

**Authors:** Ruth Hunegnaw, Anna Honko, Lingshu Wang, Derick Carr, Tamar Murray, Wei Shi, Caitlyn N. M. Dulan, Kathryn E. Foulds, Krystle N. Agans, Robert W. Cross, Joan B. Geisbert, Thomas W. Geisbert, Cheng Cheng, Aurélie Ploquin, Daphne A. Stanley, Gary J. Nabel, Nancy J. Sullivan

## Abstract

Marburg virus (MARV) causes a severe hemorrhagic fever disease in primates with mortality rates in humans up to 90%. Since 2018, MARV has been identified as a priority pathogen by the WHO, needing urgent research and development of countermeasures due to the high public health risk it poses. Recently, the first case of MARV in West Africa underscored the significant outbreak potential of this virus. The potential for cross border spread as had occurred during the Ebola 2014-2016 outbreak illustrates the critical need for Marburg vaccines. To support regulatory approval of the ChAd3-Marburg vaccine that has completed Phase I trials, we show that a non-replicating chimpanzee-derived adenovirus vector with a demonstrated safety profile in humans (ChAd3) protected against a uniformly lethal challenge with Marburg-Angola. Protective immunity was achieved within 7 days of vaccination and was maintained through one year post vaccination, antigen-specific antibodies were a significant immune correlate of protection in the acute challenge model (*p=0*.*0003*), and predictive for protection with an AUC = 0.88. These results demonstrate that a single-shot ChAd3 MARV vaccine generated a protective immune response that was both rapid and durable with a significant immune correlate of protection that will support advanced clinical development.

**One Sentence Summary:** A single-shot of non-replicating ChAd3-MARV vaccine demonstrated both rapid (within 1 week) and durable (12 months) protection against lethal Marburg virus infection in macaques.

## INTRODUCTION

Marburg virus (MARV) and Ebola virus are prototype members of the filovirus family that cause severe and lethal hemorrhagic fever disease. Marburg virus disease (MVD) outbreaks have reported case fatality rates ranging from 23-90% since its discovery in 1967. The largest MVD outbreak began in the fall of 2004 in the Uíge Province of Angola (*1*) and resulted in 252 documented cases and 227 (90%) fatalities. Most outbreaks have occurred in southern or Central Africa in the countries of Uganda, Angola, Democratic Republic of the Congo, Kenya and Zimbabwe. However, a case was recently identified in West Africa in the Guéckédou district of Guinea, the same region where the first cases of the largest Ebola outbreak were identified in 2014. This marks the first case of MVD in West Africa (*2*) and suggests that, like Ebola, there is a potential for geographic spread and this pathogen requires urgent pandemic preparedness activities to stockpile vaccines for immediate outbreak response as concluded by the WHO convened consortium of filovirus experts and vaccine stakeholders (*3*).

MVD is a zoonotic disease that can then be transmitted person-to-person via contact with contaminated blood or bodily fluids by needle-stick or exposure of mucosa, resulting in an increased risk to healthcare workers or family members who care for the infected persons (*4*). The Egyptian fruit bat (*Rousettus aegyptiacus*) is known to be one of the reservoirs for the virus (*1, 5, 6*), and a number of cases have arisen when mine workers or tourists enter caves inhabited by these bats.

Following the incubation period of 3-21 days, MVD illness begins with a generalized malaise, a high fever, and severe headache with many patients progressing to severe hemorrhagic manifestations within about a week. These symptoms include rash, conjunctival hemorrhage and bleeding from venipuncture sites (*4*). Disease progression in severe cases include renal impairment, multi-organ failure and shock. Currently, there are no approved vaccines or therapeutics for MVD which means treatment is based on supportive care measures such as oral or intravenous rehydration and management of specific symptoms.

Cynomolgus and rhesus macaques are considered the standard for modeling human filoviral disease as the clinical signs and disease progression are similar and well-characterized. Studies have shown vaccine protection in nonhuman primates (NHP), with most of the challenges limited to between 3 and 8 weeks following immunization to demonstrate protection at or near peak immune responses. Potential vaccines have included gene based approaches such as DNA or viral vectors expressing MARV glycoprotein (GP) (*7-13*), or protein based approaches such as alphavirus-vectored virus-like replicon particles (VRP) (*14*), virus-like particles (VLP) expressing MARV GP with or without NP or VP40 (*15, 16*) and recombinant protein subunit vaccines (*17*).

Recombinant adenovirus vectors (rAd) have shown protective efficacy against filoviruses either as a component of prime boost regimens, as demonstrated for Ebola virus (EBOV) showing uniform nonhuman primate (NHP) protection (*18-20*) and also as the first single-shot vaccine to demonstrate efficacy against lethal EBOV infection in macaques (*21, 22*)]. Due to the limitations of preexisting immunity to human adenovirus vectors, a replication-defective chimpanzee adenovirus was selected as an alternative vector due to the favorable safety profile and low seroprevalence in humans (*22-26*). A ChAd3-Ebola vaccine rapidly protects macaques against lethal Ebola infection after single-shot vaccination and provides durable protection through one year (*22*). The ChAd vector has been developed as a vaccine for multiple viruses and therefore has considerable clinical history in >13 Phase 1, 2 and 3 human clinical trials where it was used safely in >5,000 people, including >600 pediatric patients (*24, 26-34*). These trials demonstrated that the vector is safe and elicited antibody titers comparable to those associated with protection in NHP.

Here, we demonstrate that a single-shot ChAd3-MARV vaccine uniformly protects macaques from lethal MARV infection at both acute, and long-term memory time points. We also defined a preliminary immune correlate of protection, necessary for regulatory approval by the FDA Animal Rule. Further, ChAd3-MARV also provided rapid immunity, uniformly protecting macaques challenged as soon as 1 week after vaccination. The vaccine has been manufactured under GMP and there are ∼700 doses available for immediate outbreak response. These results in conjunction with the safety and immunogenicity demonstrated in humans highlight the potential for ChAd3-vectored single shot filovirus vaccines for use in ring-vaccination protocols (*35*) to interrupt disease transmission, while also providing significant durable immunity.

## RESULTS

### A single-shot ChAd3-MARV vaccine confers protection against acute lethal MARV challenge

Chimpanzee-derived Ad vectors are currently undergoing extensive development as vaccines in clinical trials for multiple pathogens such as Hepatitis B Virus (NCT04297917), EBOV (*23*) and SARS-CoV-2 (*36*).. For Ebola virus, ChAd3 vaccines have been shown to confer uniform protection against lethal EBOV challenge five weeks after immunization (*22*). Here, we evaluated efficacy of a replication-defective ChAd3-vectored vaccine for protection against another filovirus, MARV, in NHP. Since the ChAd3-EBOV vaccine uniformly protected NHP at a single-dose of 1 × 10^10^ particle units (PU), (*22*), the same dose was selected for ChAd3-MARV along with a second study arm in which the vaccine dose was reduced by 1-log. Groups of four cynomolgus macaques were immunized intramuscularly (i.m.) with a single inoculation of 1 × 10^10^ PU or 1 × 10^9^ PU of ChAd3 encoding MARV GP and then challenged with a uniformly lethal dose (1,000 PFU, i.m.) of MARV/Angola 5 weeks later (Fig. 1A).

**Figure 1.**
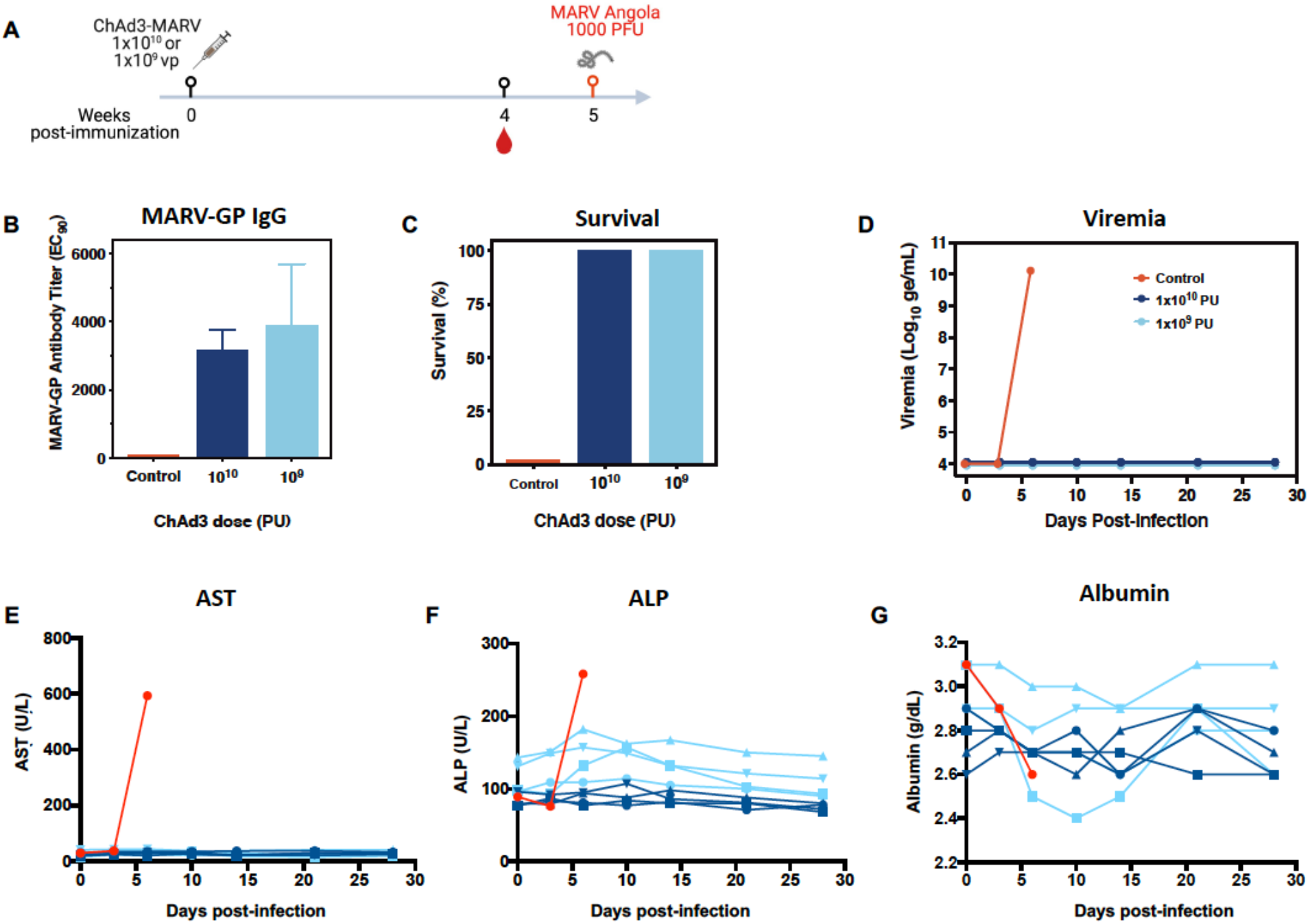
Single-shot ChAd3-MARV vaccine uniformly protects against lethal MARV/Angola challenge. (A) Vaccination scheme. Cynomolgus macaques were vaccinated with ChAd3-MARV at a dose of 1×10^10^ or 11×0^9^ PU via the i.m. route (*n* = 4 per group) or unvaccinated (*n* = 1; historical controls not shown *n* = 6). Five weeks post-vaccination, NHP were challenged with 1000 PFU MARV/Angola and followed for 28 days. (B) Plasma MARV GP–specific ELISA titers at week 4 post-vaccination (EC_90_). (C) Percent of NHP protected from death in each dose group (*n* = 4). (D) Plasma viremia assessed by quantitative reverse transcription PCR (qRT-PCR) and expressed as genome equivalents per milliliter (ge/mL). (E – G) Blood chemistry: (E) AST, aspartate transaminase; (F) ALP, alkaline phosphatase and (G) albumin levels in plasma.

MARV GP-specific antibodies were measure by ELISA one month after vaccination and showed that the 1 × 10^10^ PU and 1 × 10^9^ PU vaccine doses induced comparable titers of pre-challenge GP-specific antibodies (*p=0*.*34*) with mean EC_90_ titers of 3156 and 3884, respectively (Fig. 1B). After infectious challenge, the unvaccinated macaque succumbed to infection at day 9 post-exposure and exhibited a viral load exceeding 1 × 10^7^ genome equivalents per milliliter (ge/ml) in plasma (Fig. 1C and D). All vaccinated macaques survived the infection with no detectable viremia (Fig. 1C and D). To assess liver and kidney disease, characteristic of MVD, blood chemistry measurements were conducted for circulating levels of AST and ALP (liver) and albumin (kidney). Increases in AST and/or ALP signify inflammation or damage to liver cells leading to leakage of liver enzymes into the blood stream. On the other hand, loss of albumin, normally found at a constant level in the blood stream, can be indicative of kidney damage resulting in urinary excretion of albumin (*37*). As observed in previous NHP models for MARV infection (*38*), the unvaccinated animal showed increasing levels of AST and ALP that reached six and three times baseline levels, respectively, as well as decreased albumin in plasma by day 5, indicating organ disease in the unvaccinated control consistent with MVD (Fig. 1E to G). Animals in the 1 × 10^10^ PU ChAd3 vaccine dose group remained free from any clinical signs of disease as indicated by the maintenance of plasma liver enzyme and albumin levels within the normal range (Fig. 1E to G). At the lower ChAd3-MARV vaccine dose (1 × 10^9^ PU), a single subject showed a transient increase in ALP and a drop in albumin between days 7 and 10 post-exposure for both parameters indicating organ dysfunction, but it was transient, returning to baseline by day 21. Notably, this animal never exhibited detectible viremia or overt symptoms of disease. Together, these data demonstrate that a single-shot ChAd3-MARV vaccine can generate acute protective immunity against the lethal effects of MARV infection and a vaccine dose of 1 × 10^10^ PU protects against all signs of MVD.

### MARV GP IgG titers are an immune predictor of protection

We next performed dose-ranging studies to define the minimal protective vaccine dose and to identify a candidate immune correlate of protection under conditions of infection breakthrough. Cohorts of cynomolgus macaques were vaccinated with ChAd3-MARV at doses ranging between 1 × 10^6^ to 1 × 10^9^ PU. MARV GP-specific antibody titers were measured four weeks later (Fig. 2A). In addition to data obtained from these vaccinations, anti-MARV GP titers and percent protection data from vaccine dose groups described in Figure 1: 1 × 10^10^ (*n* = 4) and 1 × 10^9^ PU (*n* = 4) (Fig. 1B and C), were combined with data from the dose-ranging study for composite data representation across all doses in Figure 2 (Fig. 2). All 4 unvaccinated macaques succumbed after a 1000 PFU MARV/Angola challenge. Macaques in the 1 ×10^10^ and 1 ×10^9^ PU vaccine group remained viremia free (Fig. 2C), while transient viremia was observed in macaques that survived at doses of 1 ×10^8^ and lower (Fig. 2C). Average EC_90_ values across ChAd3 vaccine dose groups revealed a dose response. The average anti-GP EC_90_ titer generated in animals vaccinated at uniformly protective vaccine doses of 1 × 10^10^ (Fig. 1B, *n* = 4) and 1 × 10^9^, (Fig. 1B, *n* = 4 and Fig 2A, *n* = 4) was 4586. A reduction of vaccine dose to 1 ×10^8^ PU resulted in a decreased anti-GP EC_90_ titer to 725 (*p=0*.*014)* and infection breakthrough in 3 out of 12 macaques. Further reduction in dose to 5 × 10^7^ and 1 × 10^7^ did not result in significantly lower antibody titers, and the breakthrough rate was also maintained with 1 out of 4 macaques succumbing (Fig. 2A and B). When combined, the average EC_90_ titer generated from the 1 ×10^8^, 5 × 10^7^ and 1 × 10^7^ PU doses, where comparable titers and breakthrough were observed, was 624, significantly lower than mean titer of 4586 obtained at uniformly protective doses (*p=0*.*001*). Further dose down of the vaccine to 5 × 10^6^ PU and 1 × 10^6^ PU resulted in a combined mean titer of 53, significantly lower than previous mean titer (624) generated at higher doses (*p=0*.*003*). This reduction associated with an increase in breakthrough with 6 out of 8 macaques succumbing (Fig. 2A and B). Overall, 1 × 10^9^ PU ChAd3-MARV was the minimum uniformly protective dose in these studies and significant reduction in antibody titers was associated with increased breakthrough.

**Figure 2.**
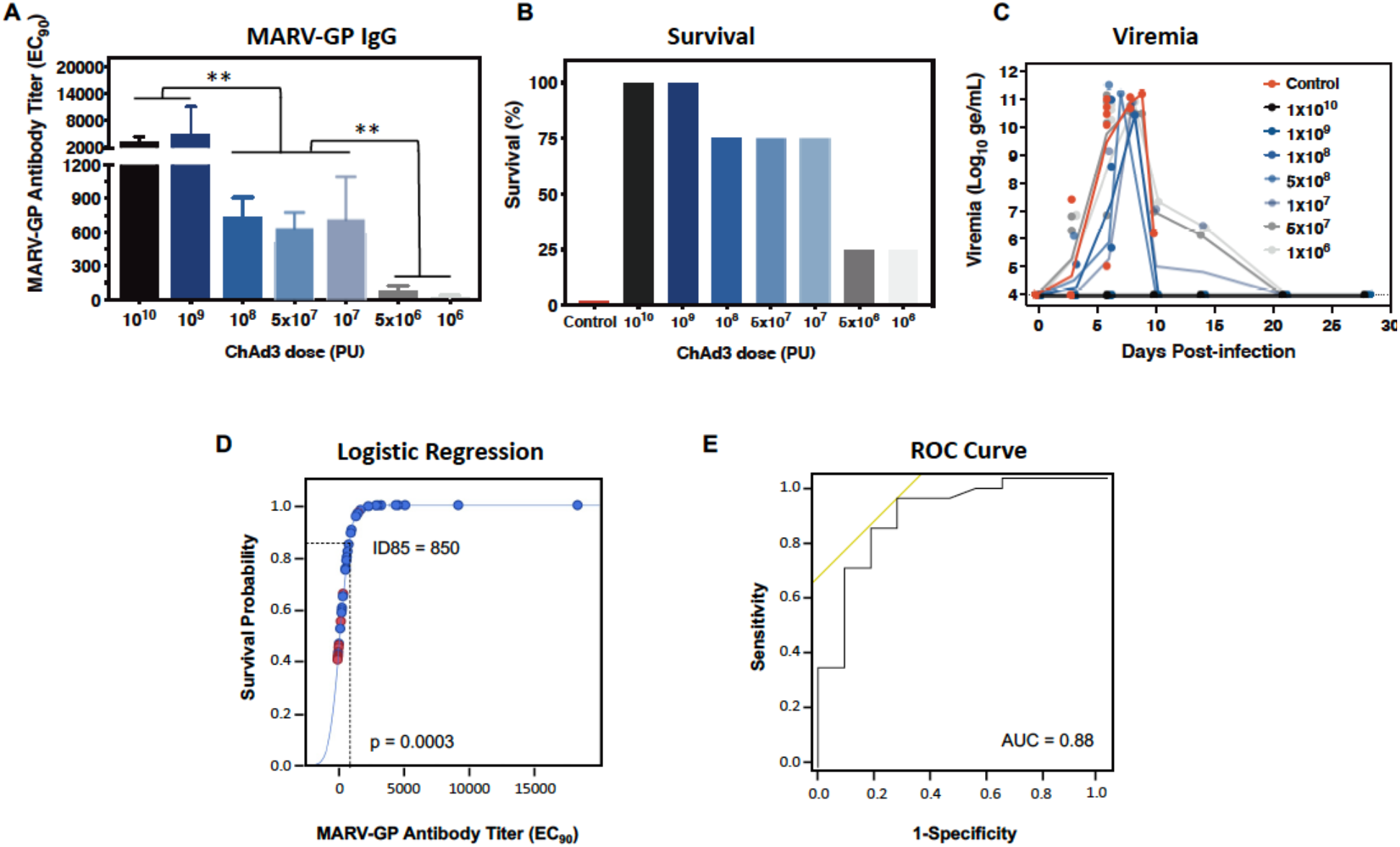
Anti-MARV GP antibodies correlate with ChAd3-MARV vaccine protection against MARV/Angola challenge. Groups of cynomolgus macaques were vaccinated with ChAd3-MARV at 1×10^9^ (*n* = 4), 1×10^8^ (*n* = 12), 5×10^7^ (*n* = 4), 1×10^7^ (*n* = 4), 5×10^6^ (*n* = 4) and 1×10^6^ (*n* = 4) via the i.m. route or unvaccinated (*n* = 4; historical controls not shown *n* = 3). Data from Fig. 1 animals (*n* = 4 per group) were included in composite data shown for vaccine dose groups 1×10^10^ and 1×10^9^ and in panels D and E analyses. Five weeks post-vaccination, NHP were challenged with 1000 PFU MARV/Angola and followed for 28 days. (A) Plasma MARV GP-specific ELISA titers at week 4 post-vaccination (EC_90_). (B) Percent of NHP protected from death in each dose group. (C) Mean plasma viremia for each vaccine dose group, as assessed by quantitative reverse transcription PCR (qRT-PCR) and expressed as genome equivalents per milliliter (ge/mL). (D) Univariate logistic regression of anti-MARV GP EC_90_ titers and survival after lethal MARV challenge. Regression was performed on ELISA IgG titers generated by ChAd3-MARV vaccination of cynomolgus macaques (*n* = 40). Y-axis illustrates survival probabilities predicted by IgG titers on a continuous scale. Blue circles, survivors; red circles, non-survivors. Dotted lines: antibody titer associated with a survival probability of 85%. The *p*-value = two-sided Likelihood ratio test (chi-square) of the slope. (E) Receiver-operating characteristic (ROC) curve of data presented in panel D; AUC, area under the curve. Statistical analysis between two groups was performed using an unpaired student’s *t*-test. (\*\**p*<0.01)

Across individual animals in these studies, near uniform protection (95%) was observed above a titer of 420, with the only exception being one animal that succumbed with a titer of 1567. Below a titer of 100, mortality was 80%, with only 2 animals out of 10 NHP surviving (Table S1). In the 100-420 titer range, both survival and mortality (22%) were observed, with the low incidence of mortality indicating that this titer range may be near the threshold titer needed for protection. In summary, these data indicated that GP-specific antibody titers associated with protection.

The associations of antibody titer with survival and the observation that serum GP-specific antibody levels are a significant predictor of vaccine-induced acute protection against EBOV (*39*), suggested antibody as a potential immune correlate of protection for the ChAd-MARV vaccine. Therefore, we tested the strength of the association between MARV GP-specific titers and survival using univariate regression analysis with IgG titer as a continuous variable, and calculated survival probabilities over the range of antibody titers observed. A significant association was found between serum anti-MARV GP titers and survival (*p=0*.*0003*) indicating that levels of anti-MARV GP positively correlate with survival (Fig. 2D). For EBOV, a GP-survival probability of 85% was benchmarked for the immune correlate (*40*). Here, we identified a MARV GP-specific titer of 850 as a correlate of 85% survival (Fig. 2D). Receiver-operating characteristic (ROC) analysis was performed to measure the sensitivity and specificity of anti-MARV GP titers in predicting survival. An area under the curve (AUC) value representing the logistic model sensitivity and specificity (where the maximum is 1.0) of 0.88 was obtained, indicating that MARV GP-specific antibody titers at 4 weeks post-vaccination provide robust discrimination between survivors and non-survivors. Therefore, the titer at 4 weeks post-vaccination is a sensitive predictor of acute protection induced by the ChAd3-MARV vaccine (Fig. 2E).

### ChAd3-MARV vaccination provides rapid protection against MARV challenge

In an outbreak setting, particularly for a virus with pandemic potential, a vaccine that confers rapid protection is ideal for ring vaccination strategies. To this end, we sought to determine how early following immunization with ChAd3-MARV, protection against MARV/Angola is conferred. Groups of 4 macaques were immunized with ChAd3-MARV at intervals of 5, 4, 3, 2 and 1 week prior to challenge. After infectious challenge with MARV/Angola the unvaccinated control exhibited a plasma viral load exceeding 1 × 10^8^ ge/mL, elevated AST and ALT and decreasing albumin before succumbing on day 9 (Fig. 3D to G). Macaques in all 5 vaccination groups were protected and remained free of plasma viremia, and levels of AST, ALT and albumin did not deviate from baseline throughout the 28 days the macaques were followed post-challenge (Fig. 3D to G). Plasma anti-MARV GP titers were assessed 1 week prior to challenge for each group (Fig. 3A). Vaccination in the 5-, 4-, and 3-week groups all resulted in mean pre-challenge GP-specific EC_90_ titers above 1100, above the 850 EC_90_ titer that predicted 85% protection (Fig. 2D). All macaques in those vaccine groups were protected from the lethal effects of MARV/Angola infection. Vaccinations at 2- and 1-week prior to challenge also uniformly protected macaques against MARV, even though mean pre-challenge GP-specific titers were 77 and 0, respectively, a range that associated with mortality when challenge was one month after vaccination, suggesting that the immune correlate of protection determined at near peak post-vaccine immunity (2-5 weeks), may not predict protection for all vaccine-challenge intervals, especially those very close to vaccination where circulating antibody responses have not yet reached peak circulating levels. Together, these data demonstrate that near immediate protection from mortality, disease, and viral replication is rapidly achieved with the single-shot ChAd3-MARVvaccine.

**Figure 3.**
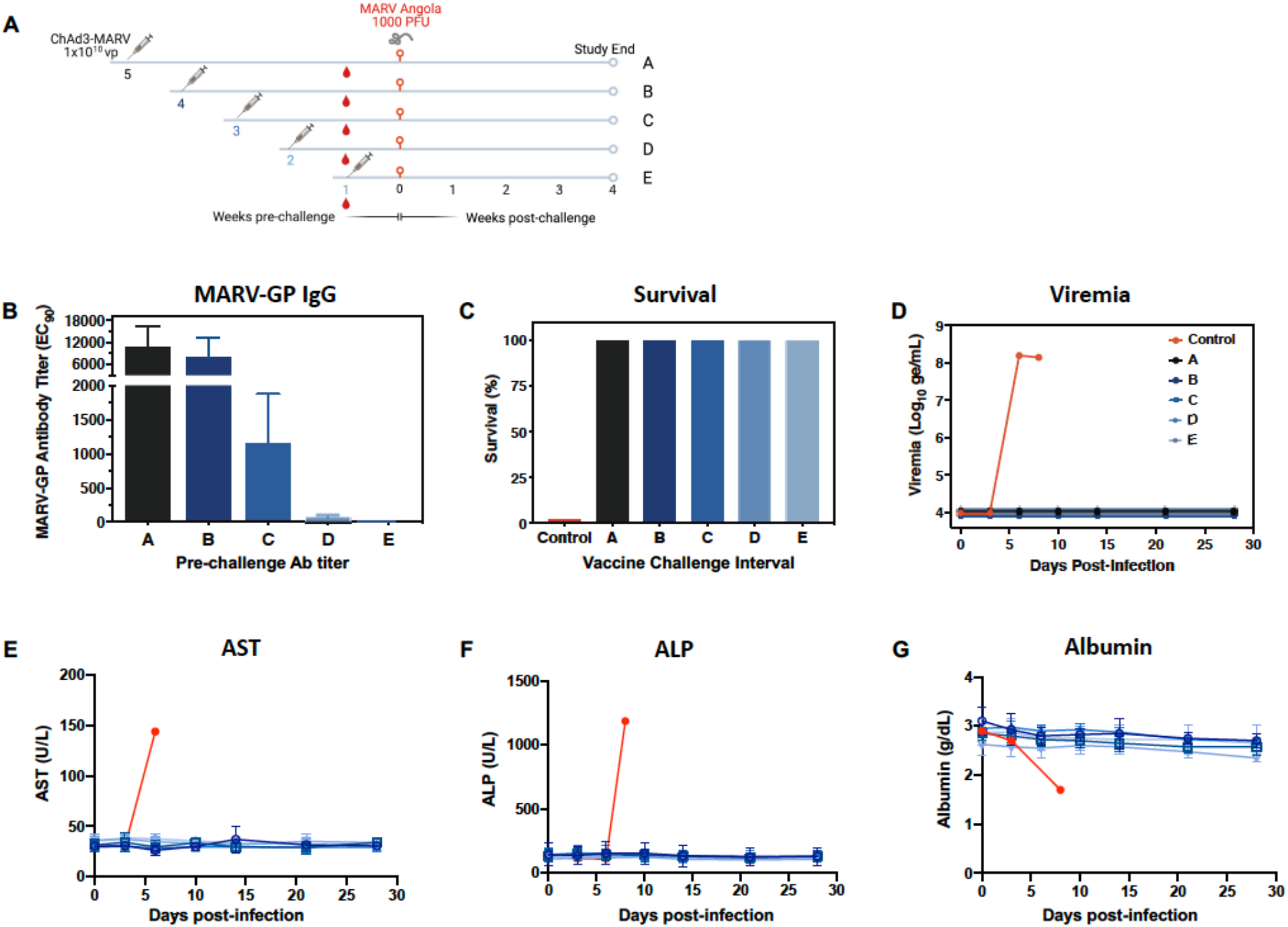
Rapid protection against MARV challenge with a single-shot ChAd3-MARV vaccine. (A) Schematic representation of vaccine groups and schedule. Groups of 4 NHP were vaccinated with ChAd3-MARV at a dose of 1 × 10^10^ PU via the i.m. route at 5, 4, 3, 2 or 1 week(s) pre-challenge or unvaccinated (*n* = 1; historical controls not shown *n* = 6). All NHP were challenged at the same time with 1000 PFU MARV/Angola and followed for 28 days. (B) Plasma MARV GP–specific ELISA titers at week 4 post-vaccination (EC_90_). (C) Percent of NHP protected from death in each vaccine group. (D) Plasma viremia for each indicated vaccine group, as assessed by quantitative reverse transcription PCR (qRT-PCR) and expressed as genome equivalents per milliliter (ge/mL). (E – G) Blood chemistry: (E) AST, aspartate transaminase; (F) ALP, alkaline phosphatase and (G) albumin levels in plasma.

### Single-shot ChAd3 vaccination provides durable protection against lethal MARV challenge

While rapid protection for ring vaccination during an outbreak response is one objective of Marburg virus vaccines, durable protection is also important for populations in filovirus endemic regions where outbreaks are becoming more periodic. Moreover, health care workers may require vaccine protection over the course of an outbreak that can last many months, so it is critical to define the durability of protective immunity. To investigate if the single-shot ChAd3-MARV vaccine can also provide extended immunologic protection, macaques (*n* = 8) were inoculated with 1 × 10^10^ PU ChAd3-MARV. Half of the animals (*n* = 4) were exposed to a lethal dose of MARV/Angola at 6 months and the remaining half at 12 months following vaccination (Fig. 4A). At one month post vaccination, the mean anti-MARV GP serum antibody titer across all eight animals was 4054 (Fig. 4A). The 6 months post-vaccination group of macaques exhibited a mean pre-challenge MARV GP-specific IgG titer of 875, a significant drop compared to the mean titer at one month post vaccination (*p=*0.019) (Fig. 4B), but above the titer that predicted 85% survival (Fig. 2D). After infectious challenge, the unvaccinated control exhibited plasma viremia that reached over 1 × 10^9^ ge/mL, as well as elevated levels of AST, ALT and diminished albumin in plasma, subsequently succumbing to the lethal effects of disease on Day 8 (Fig. 4D to G). All macaques in the six-month vaccine-challenge interval group were protected from death (Fig. 4C). In the 12-month group, macaques exhibited mean MARV GP-specific IgG titer of 446, not significantly lower than the mean titer measured at 6 months. ChAd3-MARV conferred protection in 3 out of 4 animals challenged at one year post vaccination. (Fig. 4C).

**Figure 4.**
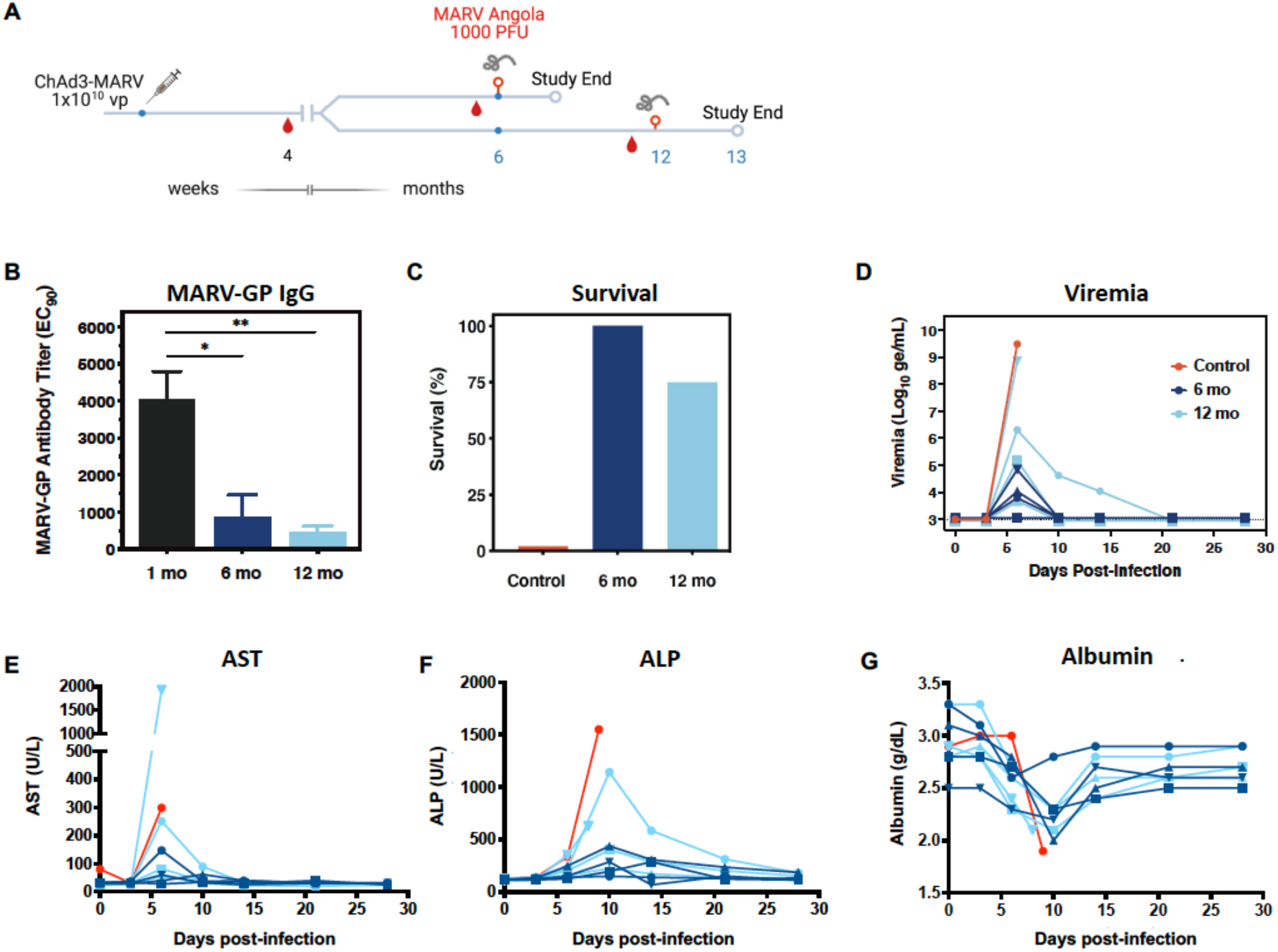
ChAd3-MARV vaccine protects against MARV challenge at durable time points. Cynomolgus macaques (*n* = 8) were immunized with ChAd3-MARV at a dose of 1×10^10^ PU via the i.m. route or unvaccinated (*n* = 1). At 6-(*n* = 4) and 12-months (*n* = 4) post-vaccination, immunized NHP were challenged with 1000 PFU MARV/Angola and followed for 28 days. (B) Plasma MARV GP–specific ELISA titers at week 4 post-vaccination (EC_90_). (C) Percent of NHP protected from death in each vaccine group. (D) Plasma viremia for each indicated vaccine group, as assessed by quantitative reverse transcription PCR (qRT-PCR) and expressed as genome equivalents per milliliter (ge/mL). (E – G) Blood chemistry: (E)AST, aspartate transaminase; (F) ALP, alkaline phosphatase and (G) albumin levels in plasma. Statistical analysis between two groups was performed using an unpaired student’s *t*-test. (\**p*<0.05, \*\**p*<0.01)

Unlike observations when challenge occurred one month after vaccination (Fig. 1), transient, low-level viremia was observed in 3 out of 4 surviving macaques from the 6-month challenge, with a peak on day 6 post-challenge but cleared by day 10 (Fig. 4D). Transient elevation in AST in one animal and ALP for a different animal between days 3 and 10 was also observed (Fig. 4E and F). In the 12-month challenge group, all 3 surviving macaques had detectable viremia, which cleared by day 10 for 2 survivors, similar to what was observed for survivors in the 6-month group. Viremia in the 3^rd^ survivor, which also had the lowest titer in the 12-month group (263), was detected until day 21 and this animal showed highest AST and ALP among all survivors (Fig. 4D to F). The animal that succumbed exhibited the second to lowest anti-GP titer (275) and the highest viremia in the group (Fig. 4D). This suggested that titers of 263 and 275 represented levels close to the threshold titer needed for durable protection. Macaques in both 6- and 12-month groups exhibited diminished plasma albumin levels post-infection that bottomed at day 10 for the protected animals before returning to baseline (Fig. 4G), indicating that impacts on kidney function were greater in animals challenged at durable time points (Fig. 4E and F). Overall, these data demonstrate that protective efficacy against lethal infection with a single-shot 1× 10^10^ PU dose of ChAd3-MARV extended up to 12 months post-vaccination. (Fig. 4C).

## DISCUSSION

Marburg virus has been named Category A priority pathogen by WHO and by the National Institutes of Allergy and Infectious Diseases (NIAID), signifying its high risk to public health (*41, 42*). When considering an ideal vaccine against Marburg virus, desirable features include a demonstrated safety profile, the induction of both rapid as well as durable protection, and a simple regimen for quick and easy deployment.

Previously, we demonstrated that a single dose of ChAd3-based EBOV vaccine administered 5 weeks prior to challenge was highly efficacious, protecting 100% of vaccinated NHPs from EVD and durable protection at one year (*22*). Using the same vector platform, we showed that a single-shot ChAd3-based MARV vaccine was protective as early as 1 week following vaccination and protection lasted for at least 1 year. We also defined a candidate immune correlate of protection which is essential for approval of a ChAd3-based MARV vaccine via the FDA Animal Rule. Demonstration of protection shortly after vaccination also highlights the quick-acting potential of the platform and suggests its suitability for deployment during an outbreak for protection of healthcare workers or in a ring-vaccination scenario.

One of the significant findings of this study is the identification of antibody titer as a correlate of vaccine induced protection against acute MARV infection. The immune correlate predicted 85% immune protection with an antibody titer of 850. This titer did not apply to immediate (1-2 week) challenge points where protection was achieved when circulating MARV GP-specific IgG was essentially undetectable. During the week following vaccination, B cells are still undergoing isotype switching from IgM to IgG/IgA (*43*). Development of hypermutated IgG antibodies specific to GP would take 10-14 days to appear in the circulation, and an average of 4 weeks to reach peak levels. In the meantime, upon antigen exposure, there is polyclonal B cell activation and differentiation into short-lived plasma cells that leads to secretion of IgM and IgG of low specificity, which has been shown to associate with initial control of infection (*44, 45*). Protection as rapidly as 1-week post-ChAd3-MARV vaccination demonstrates a robust vaccine-induced immune response that efficiently controls early viral replication.

Anti-MARV GP IgG was demonstrated for up to a year post-vaccination. Although there was a decline in circulating titer compared to 1-month post-vaccination, it was either sufficient for protection, or the anamnestic responses initiated upon re-exposure to MARV GP at challenge facilitated control of viral replication. The transient viremia and clinical signs of infection seen after challenging vaccinated macaques at memory time points suggests that there is durable antibody response that may be delayed due to the need to initiate antigen-specific memory B cell responses. The non-survivor in the 12-month challenge group had an anti-GP EC_90_ titer of 275 whereas another macaque in the same group with a titer of 263 survived. Variation in protection could be due to differences in how rapidly each NHP initiates an anamnestic response to reach the threshold of MARV GP-specific IgG required for protection. In the context of EBOV infection, robust B cell responses have been shown to lead to generation of potent antibodies that have been isolated several years after EBOV infection (*46, 47*). Further, for filoviruses, sustained anti-GP titers up to a year post-immunization with viral vectors given as a single shot have been demonstrated (*13, 23*). It is possible that vaccination with ChAd3-MARV induces similar robust B cell responses at the acute stage that is followed by contraction of the antibody response as seen with the decline in antibody titers. Upon encountering antigen during challenge, reactivation, proliferation, and differentiation of antigen-specific memory B cells can lead to production of MARV GP-specific IgG at the memory time point.

Based on these preclinical data, NIAID/VRC in collaboration with US Military HIV Research Program recently completed a Phase I trial that assessed safety, tolerability, and immunogenicity of the vaccine (NCT03475056), and ChAd3-MARV is undergoing advanced development by the Sabin Vaccine Institute,, where additional Phase I/II clinical trials will be conducted in collaboration with BARDA in filovirus-endemic populations in Africa to support regulatory approval of ChAd3-MARV GP vaccines for human use. Overall, preliminary results from the NIAID/VRC ChAd3-MARV trial reported that the vaccine was safe and resulted in no severe adverse events related to the vaccine. MARV GP-specific immune responses were measured at week 4 for both 1 × 10^10^ and 1 × 10^11^ doses and the vector was found to be immunogenic with geometric mean MARV GP-specific titers of 393 (183.8 to 838.7) and 661 (335.2 to 1302.6), respectively (NCT03475056). MARV GP-specific titer range of of 335 corresponds to a survival rate of 60-100% per the logistic regression derived from efficacy studies in NHP, indicating that a protective effect against MARV infection can be expected in humans with this vaccine regimen.

Overall, these studies support the use of ChAd3-MARV as an emergency vaccine suitable use in an outbreak setting.

## MATERIALS AND METHODS

### Construction of ChAd3 expressing MARV GP

The codon-optimized 2043bp MARV GP (Angola) sequence was excised from VRC 6712 with XbaI-BglII and cloned into pNEB Ad6 with XbaI-BamHI to make a shuttle vector pNEB Ad6 MARRV GP. The MARV GP sequence was cut out as a 3.0kb SpeI-AscI fragment and crossed over to SnaBI-linearized pChAd3(d)/SEAP to form the final plasmid ChAd3 MARV-GP.

### ChAd3 MARV GP rescue and purification

To rescue ChAd3-MARV GP vector, 293 cells were seeded at 5 × 10^6^ cells in 6-cm cell culture dishes and transfected with 3 μg of ChAd3-MARV GP vector DNA (PmeI-linearized) using lipofectamine 2000 transfection reagent as previously described (*48*). Transfected cells were collected 7-10 days post-transfection and lysed by freeze-thaw. Rescued MARV vectors were then amplified by serial passage on Procell-92 cells. The vector was purified by two rounds of standard CsCl gradient methods. Viral particles were collected from the gradient band and viral particles per ml (vp/ml) was calculated by measuring OD at 260 nM.

### Macaque immunization and infection

Groups of 2–5-year-old cynomolgus macaques (*Macaca fascicularis*) weighing between 3–5 kg were obtained from Covance for these immunization and challenge studies and were randomly assigned to each cohort. A total of 7 unvaccinated controls are included in this study. Animals were housed individually, and given enrichment regularly as recommended by the Guide for the Care and Use of Laboratory Animals. Subjects were anesthetized with ketamine prior to blood sampling or vaccination.

Cynomolgus macaques were injected intramuscularly (i.m.) in both deltoids with ChAd3 containing codon-optimized MARV glycoprotein at doses as indicated in the text and figure legends. No adverse effects of the adenovirus vaccination were observed acutely. Following immunization, all the animals were transferred to the Maximum Containment Laboratory (BSL-4) at Galveston National Laboratory for infection with MARV, and remained there through study completion. Surviving animals were followed for at least 4 wk post-exposure.

### Filovirus Challenge

Marburg virus Angola isolate 200501379 (MARV/Ang) was isolated from an 8-month-old female patient in Uige, Angola from serum collected on day 17 post onset, 1 day before death. Viral challenge material was from the second Vero E6 (ATCC CRL-1586) cell passage of MARV/Angola isolate 200501379, CDC 810820 (Passage 1 material). Cell supernatants were subsequently harvested at day 6 post infection and stored at −80°C until used for challenge. No mycoplasma or endotoxin were detectable (< 0.5 EU/ml).

NHP were transferred to UTMB and challenged at various intervals following immunization (ranging from 1 week to 12 months). Macaques were challenged by intramuscular injection with a lethal dose (target 1,000 PFU; actual titers ranged from 750 – 1413 PFU with a median of 1213 PFU) of MARV/Ang. UTMB study staff was blinded as to the experimental group of each animal. Physical exams were conducted, and blood was collected for chemistry, hematology assessment as well as ELISAs and viremia by qRT-PCR. NHP were monitored daily and scored for disease progression and moribund animals were humanely euthanized per UTMB IACUC-approved protocol.

Clinical observations prior to the onset of clinical signs were performed at least twice per day by study staff or animal caretakers. After the onset of clinical signs (responsiveness, altered posture and appearance, presence of rash, bleeding, gastrointestinal signs, food consumption altered, edema, respiration rate altered, exudates from any orifice, and/or altered neurological function), animals were observed more than twice daily by study staff. When macaques were anesthetized for blood collection or physical exams, body temperatures were taken using a rectal thermometer.

### Hematology and Serum Chemistry

Complete blood counts were assessed using a hematology analyzer (Beckman Coulter, Brea, CA, USA) in accordance with manufacturer guidelines. Serum chemistry analysis was obtained from whole-blood samples collected in sterile red-top tubes and processed to serum. Serum samples were tested for albumin (ALB), alanine aminotransferase (ALT), aspartate aminotransferase (AST), alkaline phosphatase (ALP), amylase (AMY), blood urea nitrogen (BUN), calcium (CA), creatinine (CRE), glucose (GLU), gamma-glutamyl transferase (GGT) and C-reactive protein (CRP) by using a Biochemistry Panel Plus disc on a Piccolo point-of-care analyzer (Abaxis, Inc., Union City, CA, USA) according to the manufacturer’s guidelines.

### Viremia by quantitative real-time RT-PCR

RNA was extracted using a Qiagen Viral RNA Mini kit (Qiagen Mississauga, ON, Canada). AVL-treated whole blood was first lysed using Qiagen Qiashredder tubes to liberate intracellular RNA. RNA was extracted according to the manufacturer recommendations and treated with DNase.

One-Step Probe RT-qPCR kits (Qiagen) and CFX96 system/software (BioRad) were used to determine viral copies in samples. To detect MARV RNA, we targeted the MARV NP gene with primer pairs and a 6-carboxyfluorescein (6FAM)–5′-CCCATAAGGTCACCCTCTT-3′–6 carboxytetramethylrhodamine (TAMRA) probe. Thermocycler run settings were 50 °C for 10 min; 95 °C for 10 s; and 40 cycles of 95 °C for 10 s plus 59 °C for 30 s. Integrated DNA Technologies synthesized all primers and Life Technologies customized probes. Viremia is expressed as log_10_ genome equivalents using an RNA genomic standard curve with a limit of detection for this assay of either 1E+3 or 1E+04 genome copies/mL based on the RNA standard.

### Anti-Marburg GP IgG ELISA

Methods for the GP IgG ELISA have been described previously (*49*). Briefly, Nunc-Immuno MaxiSorp™ plates (Nunc, Rochester, NY) were coated with 10ug/ml of lectin Galanthus nivalis (Sigma-Aldrich, St Louis, MO) at 4°C overnight and then blocked with 10% fetal calf serum at 4°C overnight followed by six washes with PBS containing 0.2% Tween 20 (Sigma-Aldrich). Prepared lectin plates were then incubated at 4°C overnight with a transmembrane deleted form of the MARV GP Popp strain (Lake Victoria MARV, strain Popp 1967, UniProtKB - P35254), washed, and incubated with serial dilutions of 1:50-1:50,000 of subject sera or plasma. Bound IgG was detected using goat anti-human IgG (Southern Biotech, Birmingham, AL) conjugated to horseradish peroxidase and Sigma Fast o-phenylenediamine dihydrochloride (Sigma-Aldrich). The optical density was determined at 450nm using a Victor X3 plate reader (Perkin Elmer, Waltham, MA). Background-subtracted ELISA titers are expressed as reciprocal optical density values, representing the dilution at which there is a 90% decrease in antigen binding (EC_90_).

### Statistical analyses

GraphPad Prism 9 and JMP 16 softwares were used to prepare figures and conduct statistical analyses. Averaged data values are presented as mean ± SEM. Comparisons of anti-Marburg GP antibody titers (EC_90_) were done using Student’s two-tailed *t*-test.

## Supplementary Materials

Fig. S1. Additional hematologic and blood chemistry data corresponding to main Fig. 1.

Fig. S2. Additional hematologic and blood chemistry data corresponding to main Fig. 3. Fig. S3. Additional hematologic and blood chemistry data corresponding to main Fig. 4.

Table S1. MARV GP antibody titers measured at 4 weeks post-immunization with ChAd3-MARV.

## Acknowledgments

This work was supported by the Intramural Research Program of the Vaccine Research Center, the National Institute of Allergy and Infectious Diseases, and the National Institutes of Health. We thank John Mascola for advice, Qiong Chen for technical assistance, Viktoriya Borisevich for assisting with clinical pathology, Karla Fenton for assisting with necropsies, Chad Mire and Daniel Deer for assisting with animal studies.

## Author contributions

Conceptualization: NJS, GJN Methodology: NJS

Formal Analysis: RH, AH, DAS, AP

Investigation: LW, DC. TM, WS, CM, KEF, KNA, RWC, JBG, CC, DAS

Project administration: TWG, NJS Supervision: NJS, GJN

Writing – original draft: RH, AH

Writing – review & editing: RH, AH, NJS

