## Supplemental Data for "Rapid and Durable Protection Against Marburg Virus with a Single-Shot ChAd3-MARV GP Vaccine"

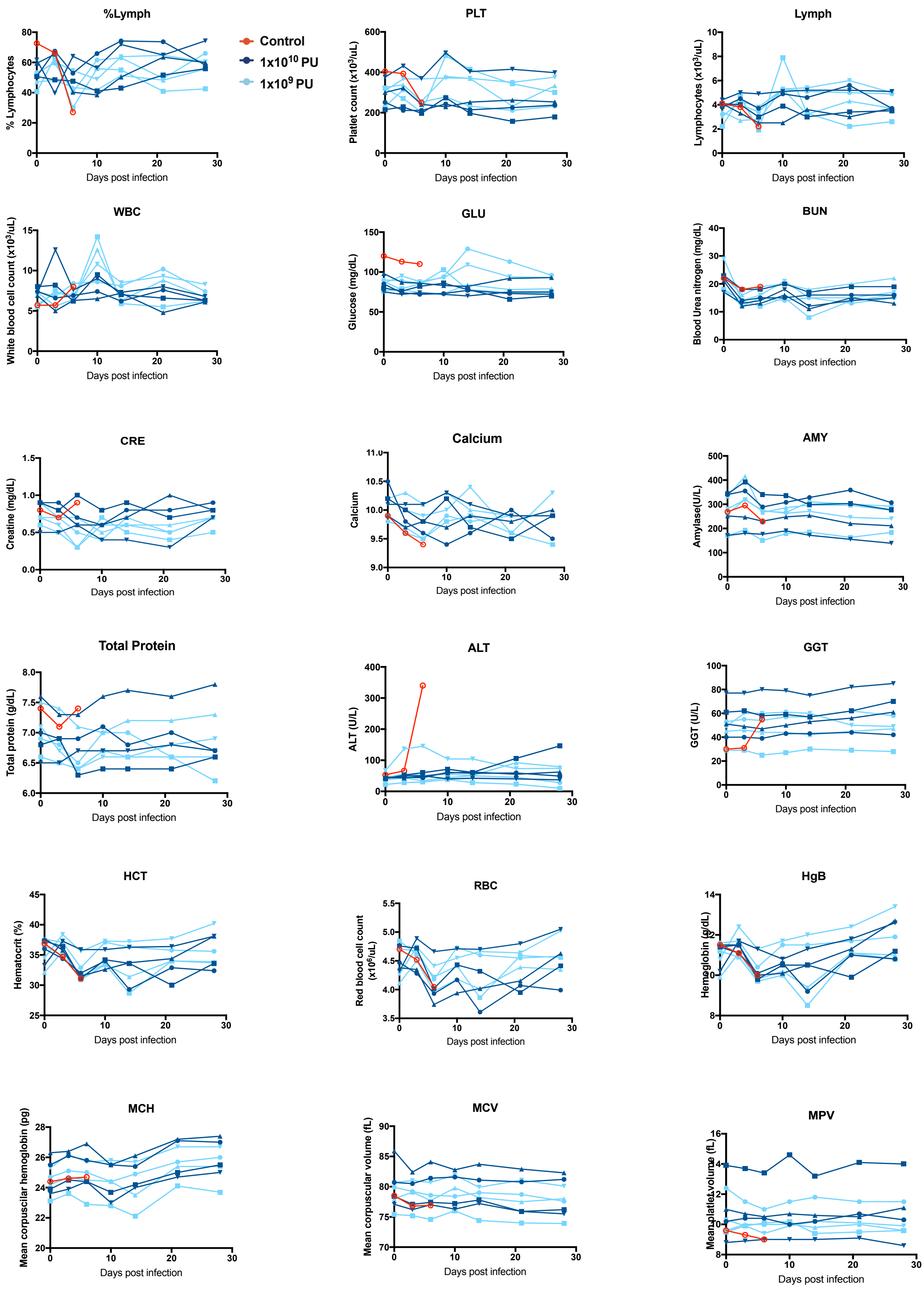
%Lymph
% Lymphocytes
Days post infection
Control
1x10$^{10}$ PU
1x10$^{9}$ PU
PLT
Platelet count (x10$^3$/uL)
Lymph
Lymphocytes (x10$^3$/uL)
WBC
White blood cell count (x10$^3$/uL)
GLU
Glucose (mg/dL)
BUN
Blood Urea nitrogen (mg/dL)
CRE
Creatine (mg/dL)
Calcium
AMY
Amylase(U/L)
Total Protein
Total protein (g/dL)
ALT
ALT (U/L)
GGT
GGT (U/L)
HCT
Hematocrit (%)
RBC
Red blood cell count (x10$^6$/uL)
HgB
Hemoglobin (g/dL)
MCH
Mean corpuscilar hemoglobin (pg)
MCV
Mean corpuscular volume (fL)
MPV
Mean platelet volume (fL)

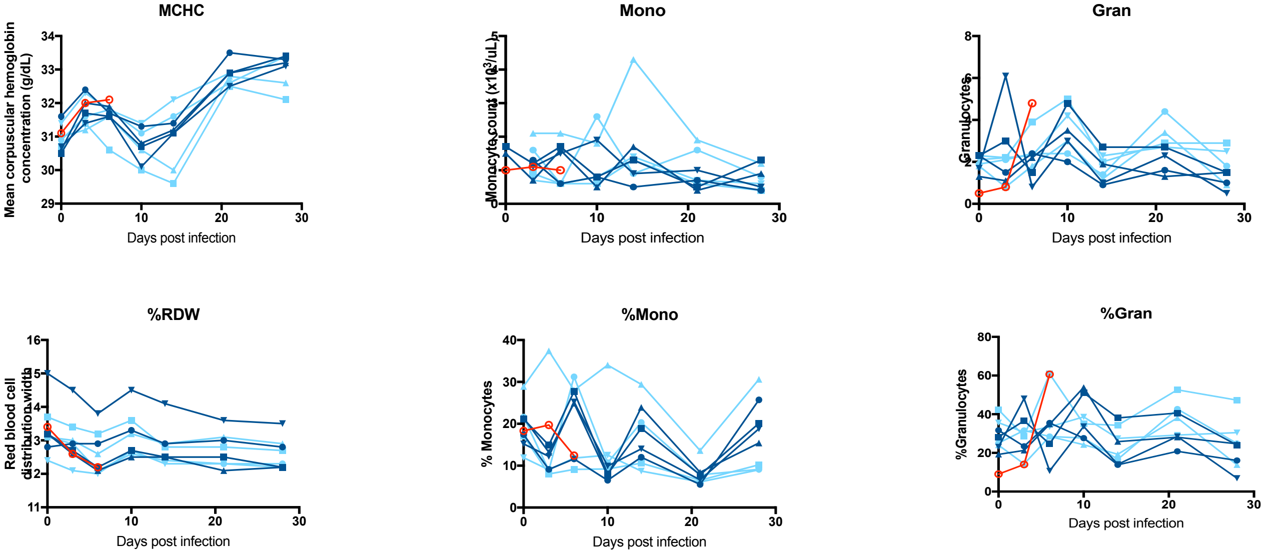


**Figure S1**. Additional hematology and blood chemistry data corresponding to Main Figure 1. Cynomolgus macaques were vaccinated with ChAd3-MARV at a dose of $1x10^{10}$ or $11x0^{9}$ PU via the i.m. route (*n* = 4 per group) or unvaccinated (*n* = 1; historical controls not shown *n* = 6). Five weeks post-vaccination, NHP were challenged with 1000 PFU MARV/Angola and followed for 28 days.

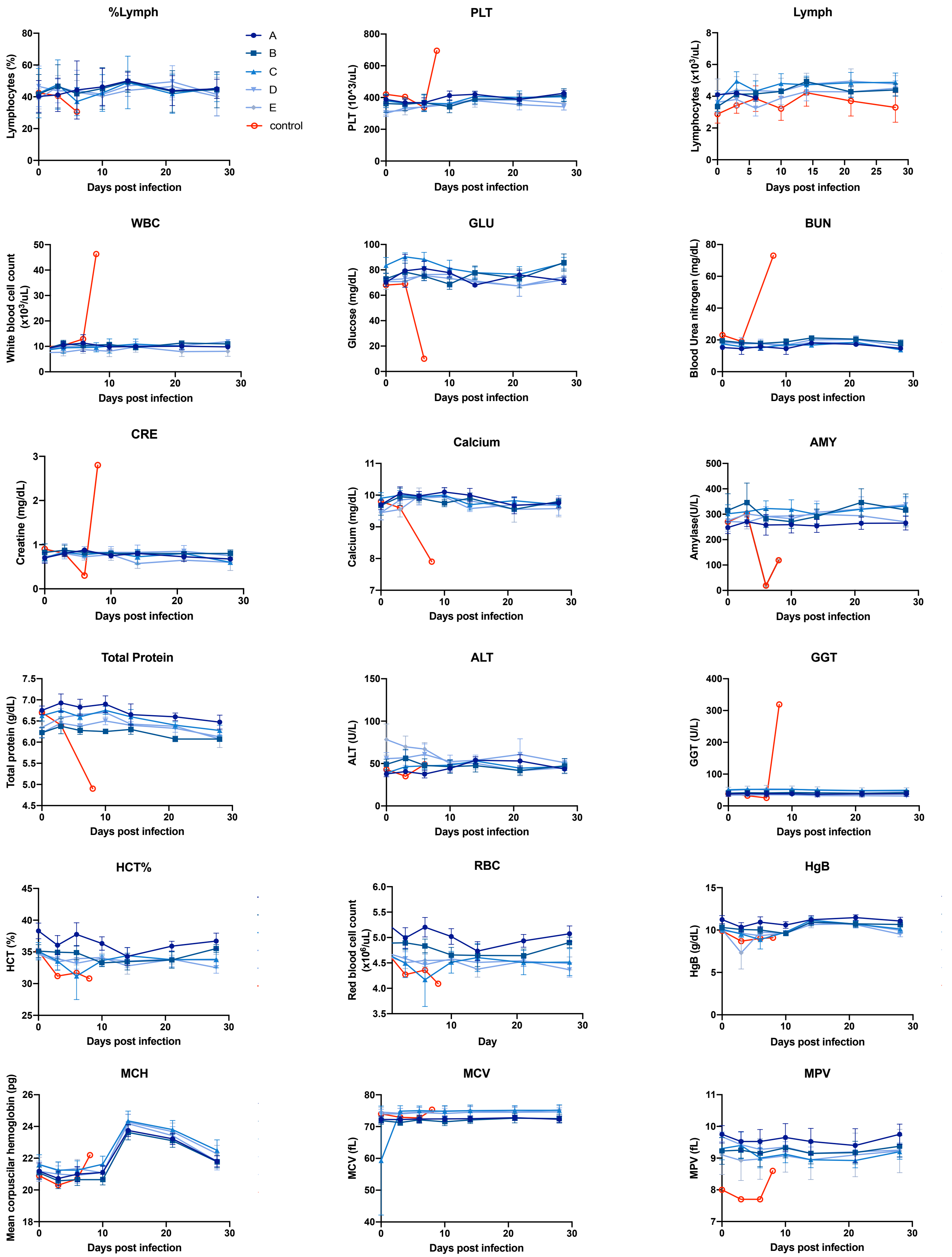
%Lymph
Lymphocytes (%)
Days post infection
A
B
C
D
E
control
PLT
PLT (10^3/uL)
Days post infection
Lymph
Lymphocytes (x10$^3$/uL)
Days post infection
WBC
White blood cell count (x10$^3$/uL)
Days post infection
GLU
Glucose (mg/dL)
Days post infection
BUN
Blood Urea nitrogen (mg/dL)
Days post infection
CRE
Creatine (mg/dL)
Days post infection
Calcium
Calcium (mg/dL)
Days post infection
AMY
Amylase(U/L)
Days post infection
Total Protein
Total protein (g/dL)
Days post infection
ALT
ALT (U/L)
Days post infection
GGT
GGT (U/L)
Days post infection
HCT%
HCT (%)
Days post infection
RBC
Red blood cell count (x10$^6$/uL)
Day
HgB
HgB (g/dL)
Days post infection
MCH
Mean corpuscular hemoglobin (pg)
Days post infection
MCV
MCV (fL)
Days post infection
MPV
MPV (fL)
Days post infection

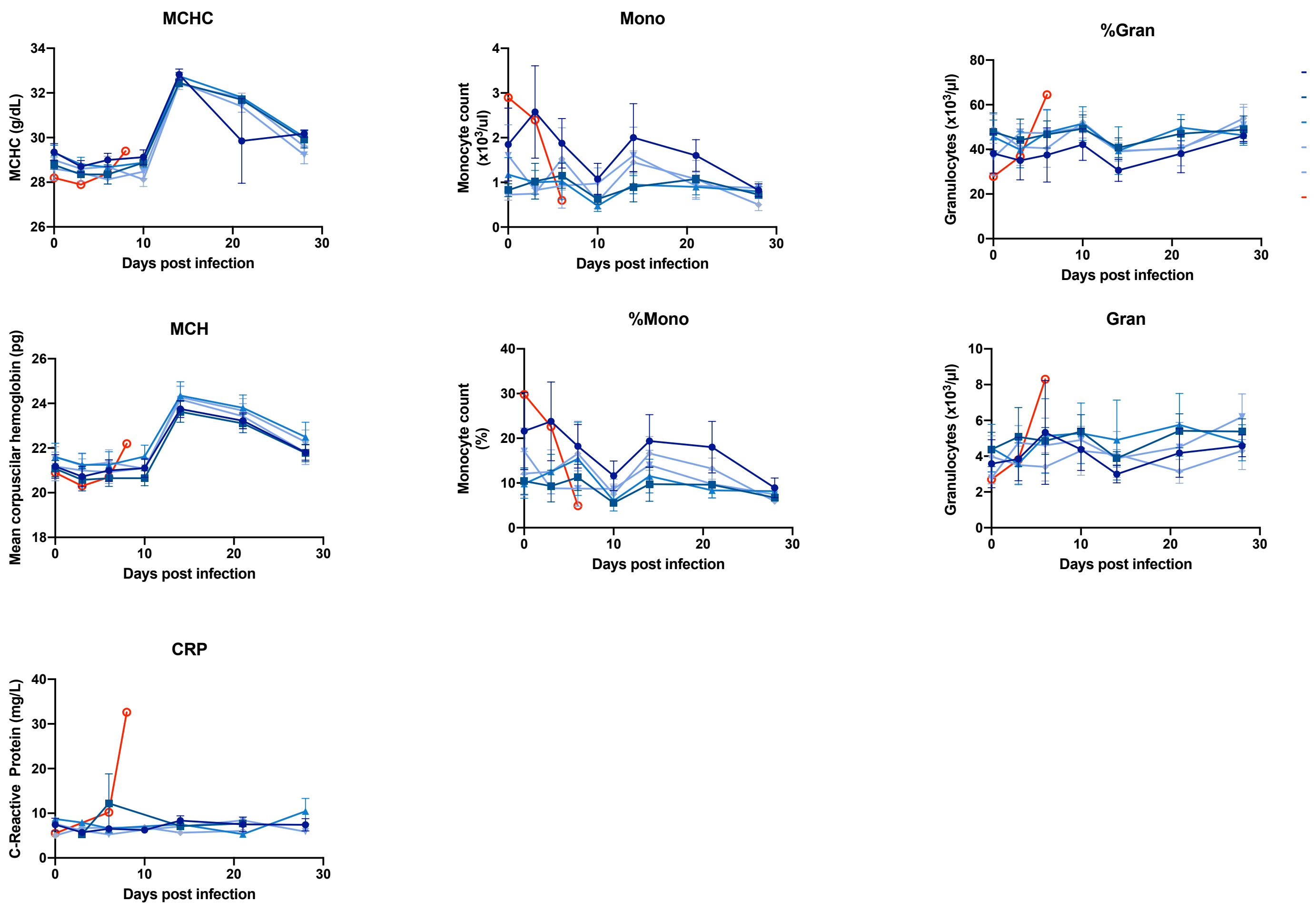


**Figure S2**. Additional hematology and blood chemistry data corresponding to Main Figure 3. Groups of 4 NHP were vaccinated with ChAd3-MARV at a dose of 1 x $10^{10}$ PU via the i.m. route at 5 (A), 4 (B), 3 (C), 2 (D) or 1 (E) week(s) pre-challenge or unvaccinated (*n* = 1; historical controls not shown *n* = 6). All NHP were challenged at the same time with 1000 PFU MARV/Angola and followed for 28 days.

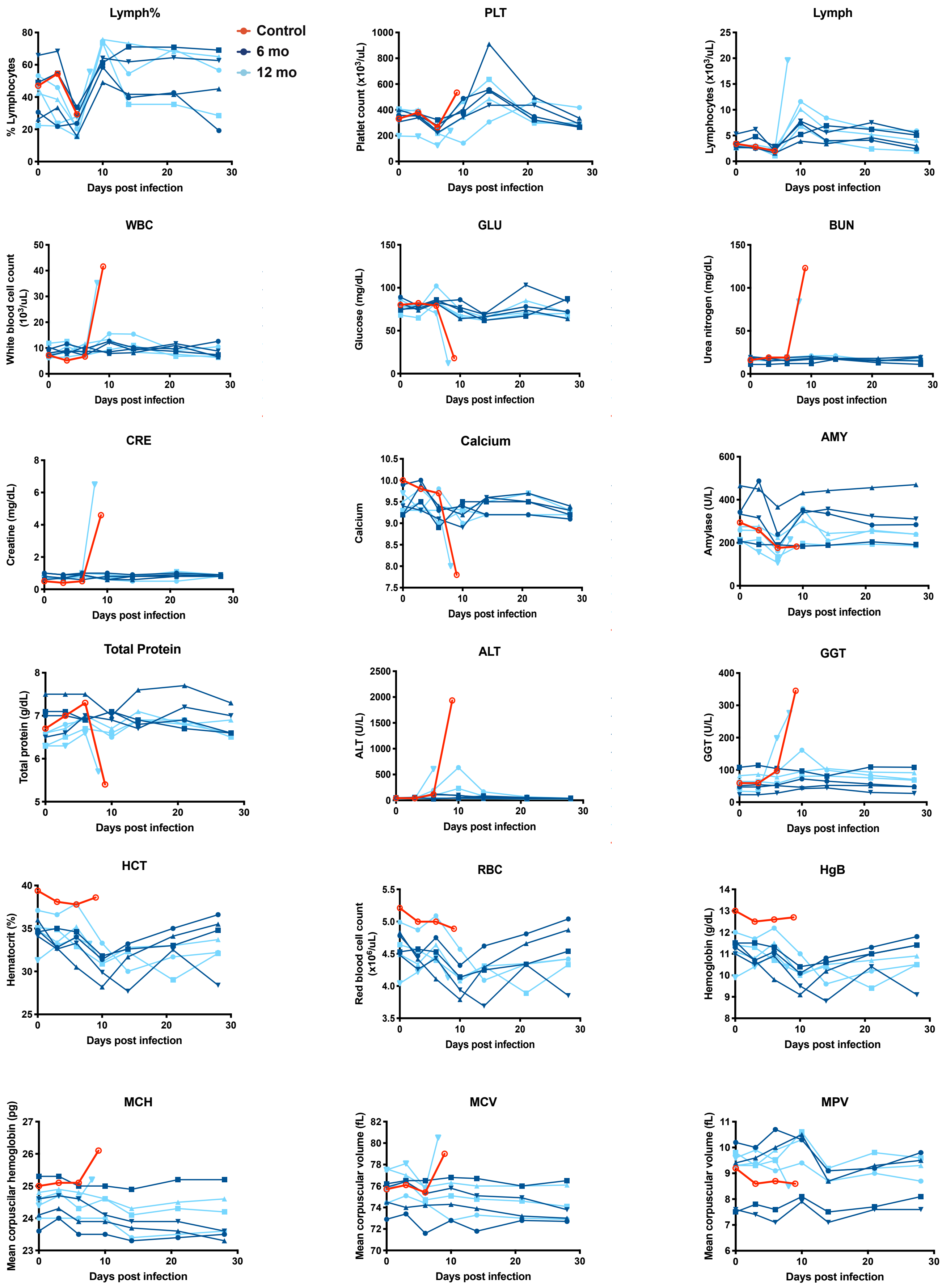

Lymph%
Control
6 mo
12 mo
% Lymphocytes
Days post infection
PLT
Platelet count (x10³/uL)
Days post infection
Lymph
Lymphocytes (x10³/uL)
Days post infection
WBC
White blood cell count (10³/uL)
Days post infection
GLU
Glucose (mg/dL)
Days post infection
BUN
Urea nitrogen (mg/dL)
Days post infection
CRE
Creatine (mg/dL)
Days post infection
Calcium
Calcium
Days post infection
AMY
Amylase (U/L)
Days post infection
Total Protein
Total protein (g/dL)
Days post infection
ALT
ALT (U/L)
Days post infection
GGT
GGT (U/L)
Days post infection
HCT
Hematocrit (%)
Days post infection
RBC
Red blood cell count (x10⁶/uL)
Days post infection
HgB
Hemoglobin (g/dL)
Days post infection
MCH
Mean corpuscular hemoglobin (pg)
Days post infection
MCV
Mean corpuscular volume (fL)
Days post infection
MPV
Mean corpuscular volume (fL)
Days post infection

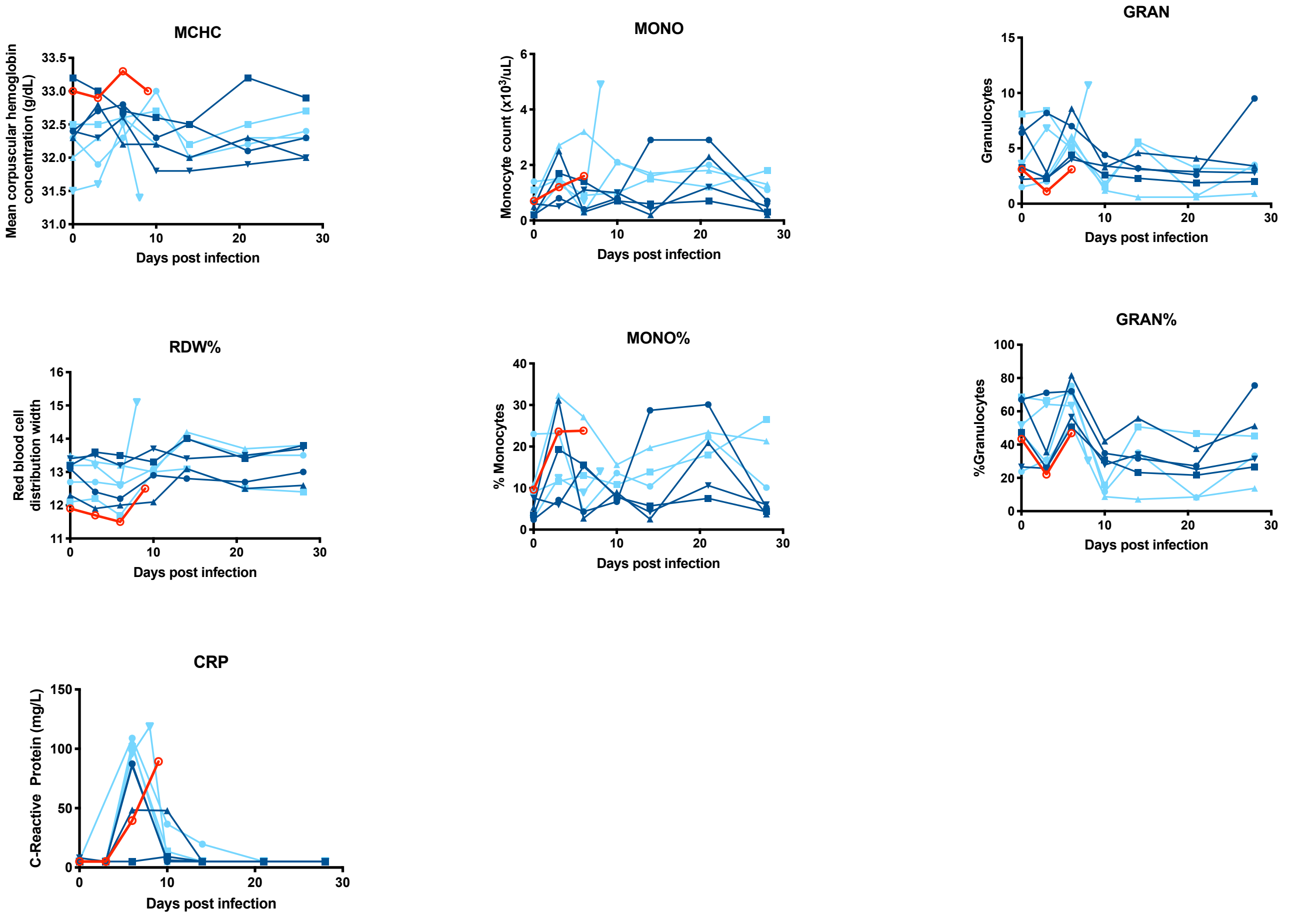


**Figure S3**. Additional hematology and blood chemistry data corresponding to Main Figure 4. Cynomolgus macaques (*n* = 8) were immunized with ChAd3-MARV at a dose of $1x10^{10}$ PU via the i.m. route or unvaccinated (*n* = 1). At 6- (*n* = 4) and 12-months (*n* = 4) post-vaccination, immunized NHP were challenged with 1000 PFU MARV/Angola and followed for 28 days.

**Table S1**. MARV GP antibody titers measured at 4 weeks post-immunization with ChAd3-MARV.

| NHP # | Titers |
|---|---|
| 1 | 18327 |
| 2 | 9167 |
| 3 | 5080 |
| 4 | 4542 |
| 5 | 4376 |
| 6 | 3341 |
| 7 | 3086 |
| 8 | 2945 |
| 9 | 2363 |
| 10 | 1764 |
| 11 | 1567 |
| 12 | 1532 |
| 13 | 1403 |
| 14 | 1063 |
| 15 | 1009 |
| 16 | 851 |
| 17 | 772 |
| 18 | 715 |
| 19 | 682 |
| 20 | 611 |
| 21 | 601 |
| 22 | 420 |
| 23 | 420 |
| 24 | 403 |
| 25 | 328 |
| 26 | 311 |
| 27 | 295 |
| 28 | 241 |
| 29 | 195 |
| 30 | 102 |
| 31 | 93 |
| 32 | 78 |
| 33 | 50 |
| 34 | 38 |
| 35 | 31 |
| 36 | 27 |
| 37 | 24.7 |
| 38 | 16 |
| 39 | 6 |
| 40 | 0 |

Red shaded cells indicate NHPs that succumbed
